# DP/MM: A Hybrid Model for Zinc-Protein Interactions in Molecular Dynamics

**DOI:** 10.1101/2023.09.29.560253

**Authors:** Ye Ding, Jing Huang

**Author notes:** Electronic Supplementary Information (ESI) available: [details of any supplementary information available should be included here]. See DOI: 00.0000/00000000.

## Abstract

Zinc-containing proteins are essential to a variety of biological processes, yet accurately modeling them using classical force fields is hindered by complicated polarization and charge transfer effects. This study introduces DP/MM, a hybrid force field model that combines *ab initio* accuracy with MM-level efficiency for modeling zinc-protein interactions. The DP/MM scheme utilizes a deep potential model to correct the atomic forces of zinc ions and their coordinated atoms, elevating them from MM to QM levels of accuracy. The model is trained on the difference in atomic forces between MM and QM calculations across diverse zinc coordination groups. Simulations on a variety of zinc-containing proteins demonstrate that DP/MM faithfully reproduces their coordination geometry and structural characteristics, for example, the tetrahedral coordination structures for the *Cys*^4^ and the *Cys*^3^*His*^1^ groups. Furthermore, DP/MM is capable of handling exchangeable water molecules in the zinc coordination environment. With its unique blend of accuracy, efficiency, flexibility, and transferability, DP/MM not only serves as a valuable tool for studying zinc-containing proteins but also represents a pioneering approach that augments the growing landscape of machine learning potentials in molecular modeling.

## 1 Introduction

Metal ions are crucial for the biological function of many proteins ^1^. These metalloproteins need the binding of one or more metal ions for their activity in biological processes. The intriguing interplay between metal ions and protein functions has stimulated intense research focus on elucidating the structure-dynamics-function relationship of these metalloproteins, employing both experimental and theoretical approaches ^2–8^. Zinc-containing proteins constitute a large fraction of metalloproteins found in the human body ^9^. Zinc ions (Zn^2+^) either directly participate in the catalytic process of enzymes, or help stabilize essential biomolecular structures ^10,11^. With their flexible electronic structure, zinc ions can form a variety of coordination complexes with protein residues and thus play vital roles in numerous biological processes, such as transcription involving zinc finger proteins ^10,12^, alcohol metabolism through the alcohol dehydrogenase (ALDH) protein family ^11,13,14^, and peptide bond hydrolysis with the carboxypeptidase (CP) family ^10,14,15^. While molecular dynamics (MD) simulation serves as an important tool in dissecting the conformational dynamics of biological macromolecules ^16,17^, it faces significant challenges when simulating zinc-containing proteins ^18^.

The accuracy of the force fields (FFs) and the adequate sampling of conformational space are generally recognized as the two crucial factors for MD simulations ^19^. FFs depict the interactions between all atoms within the simulation system and, when modeled accurately, allow for meaningful comparisons between simulation and experimental observables ^20^. On the other hand, the ability of MD simulations to efficiently and adequately explore the relevant conformational space is tightly bound to the computational efficiency of force evaluations associated with a given FF. This necessitates a delicate balance between accuracy and efficiency for FF development, which is especially difficult for simulating zinc-containing proteins ^18^. The commonly used bonded and non-bonded terms in classical FFs fall short in describing the interactions between zinc ions and amino acids which involve complicated polarization and charge transfer effects ^21–23^. While the non-bonded parameters in both pair-wised Lennard-Jones (LJ) potential and Coulomb potential terms have been fine-tuned for interactions between zinc ions and amino acids ^24–26^, it’s often observed in MD simulations that the coordination groups for zinc-containing proteins are distorted and deformed ^27–29^.

Numerous efforts have been made to improve the accuracy of modeling zinc-proteins interactions with classical FFs. It is common practice to employ harmonic bond restraints in zinc-protein interactions to ensure the maintenance of correct coordination numbers (CNs) during simulations. This means incorporating additional bonded interactions in the system’s Hamiltonian. An alternative to the bonded model is the valence bond model ^30–32^, which is based on the concept of hybridization orbitals. Modifications can also be made to the non-bonded terms, including both LJ ^33,34^ and Coulomb terms ^35^, to implicitly describe the polarization and charge transfer effects in zinc-protein interactions. Alternative approaches to account for the polarization effect include distributing the charges by introducing dummy atoms ^18,36–38^ or employing polarizable FF models such as the induced dipole model ^39–41^ and the classical Drude oscillator model ^42^. Furthermore, it’s possible to construct an accurate Hamiltonian to study a specific system. For instance, Li et al detailed the zinc-coupled folding pathway for a zinc-finger protein by dynamically including the charge transfer effect in their simulations ^43,44^. The introduction of these complex interaction terms, however, inevitably limits the transferability of these FF models across diverse coordination environments.

The hybrid quantum mechanics/molecular mechanics (QM/MM) method is widely adopted for simulating processes that are challenging to model using classical FFs ^45,46^. The QM/MM method partitions the simulation system into a QM region and an MM region, which are evaluated with QM and MM FFs, respectively. When applied to metalloproteins, the QM region typically includes the metal ions and their coordinating ligands, thereby enabling a relatively accurate modeling of the metal-protein interactions ^47–49^. Nevertheless, QM/MM models also face considerable challenges ^50^. The expensive cost of QM calculations severely limits the accessible timescale of QM/MM simulations to several orders of magnitude lower than standard MD simulations. The partition of QM and MM regions, as well as the electrostatic embedding schemes, are highly system-dependent, which reduces the adaptability of QM/MM applications ^49,51,52^. In general, the QM and the MM regions are fixed throughout the simulations, which makes QM/MM particularly difficult to apply to zinc-protein interactions, where water molecules are often involved in the zinc coordination and thus need to be dynamically exchangeable during the simulations.

In recent years, machine learning potentials (MLPs) have emerged as a compelling alternative to QM methods due to their efficiency, accuracy and transferability ^53–59^. MLPs are constructed using flexible functions such as neural networks or kernel functions and optimized by fitting them with the physical properties of systems derived from QM calculations. Once properly trained, MLPs can reproduce QM-level accuracy for properties such as energies and forces at significantly reduced computational costs. Furthermore, most MLPs decompose the system’s potential energy into the sum of atomic contributions, following the pioneering work of Behler and Parrinello ^53^, and thus scale linearly with system size. Each atomic contribution is a function of atomic environment vectors, which comprise atomic coordinates and attributes. The definitions of these atomic environment vectors are what distinguish various MLPs.For example, the ANI-1 and ANI-2 series of MLPs were developed based on the atomic environment vectors of two-body and three-body terms with atomic-centered symmetry functions ^54,60^. Deep Potential (DP) ^56^ and SchNet ^57^ model the atomic environment vectors using embedding layers and message-passing neural networks, respectively. Meanwhile, Gaussian approximation potential (GAP) and Bayesian force field employ kernel functions to construct the atomic environment ^55,58^. One of the prevailing challenges when applying MLPs to study biomolecular systems is the proper treatment of long-range non-bonded interactions.

In this work, we leverage MLPs (specifically, the Deep Potential model) to develop a novel DP/MM strategy for accurately modeling zinc-protein interactions on a timescale comparable with classical MM simulations. Unlike the QM/MM approach, which divides the simulation system into separate QM and MM regions, the hybrid DP/MM model incorporates the DP as a correction term that provides additional atomic forces to supplement the MM forces for a specific list of atoms in the system. The entire system is also evaluated using a conventional FF uniformly during the simulations. The added DP term bridges the gap between the higher-level QM and the MM forces on zinc and zinc-coordinated atoms, which can be considered as a Δ term ^61–63^. In DP/MM simulations, zinc ions and their associated ligand atoms included in the DP description are selected adaptively. Simulations of zinc-containing protein systems, including zinc finger domains and carbonic anhydrases, indicate that this hybrid DP/MM model can describe zinc-protein interactions with QM-level accuracy and MM-level efficiency.

The manuscript is organized as follows. After providing an overview of the DP model and the DP/MM strategy, we elaborate on the training and simulation settings for DP/MM in the “Methods” section. The “Results” section includes the results of DP/MM training and simulations. In particular, various zinc-containing proteins with distinct coordination groups were utilized to evaluate the performance of DP/MM. Furthermore, to investigate the zinc-water interactions in DP/MM, we also include simulation results of a zinc ion in an aqueous solution. We conclude the manuscript with a discussion of the advantages and limitations of the proposed DP/MM scheme.

## 2 Methods

### 2.1 Deep Potential Model

The Deep Potential model offers a novel way for the accurate simulation of atomic interactions, achieving QM accuracy at a substantially reduced computational cost compared to QM methods. A well-designed neural network that preserves symmetry and automatically learns atomic environment vectors is the key to the DP model. The system’s potential energy in the DP model is expressed as the sum of individual atomic energies (Eq. 1). Specifically, a neural network *NN*_fitting_ takes as input the atomic environment vectors *𝒟*_*i*_ to output the atomic energy *E*_*i*_. The *𝒟*_*i*_ vectors, in turn, are derived from a rotation-invariant embedding network *NN*_embedding_ using the relative displacement vectors 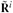 between the *i*-th atom and its neighboring *j*-th atom (Eq. 2). The fitting networks are distinguished by the element type of the central atom *i*, while variations in the embedding networks arise based on the atom pair type between the central atom *i* and its neighbor *j*.

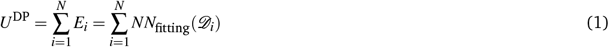

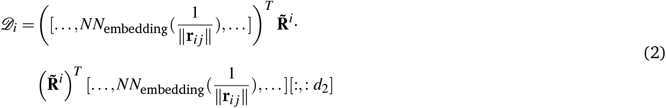

The DP model has found extensive applications in modeling various systems, such as the water phase diagram ^64^, the water– TiO2 interface ^65^, and high–entropy materials ^66^, demonstrating substantial benefits in both accuracy and efficiency. Nevertheless, the application of DP to biomolecular systems, which are more dynamic and inhomogeneous, remains limited. This is largely due to the absence of an explicit description of long-range effects in the DP architecture, a limitation shared by the majority of MLPs.

### 2.2 DP/MM Model

To extend the applicability of the DP model for simulating zinc-containing proteins, we proposed a novel DP/MM strategy for zinc-protein interactions with *ab initio* accuracy, enhanced flexibility, and reduced computational cost. The hybrid DP/MM force field model integrates DP and MM through a simplified approach, in which the entire system is governed by MM while dynamically changing regions of interest are described by additional DP models. This strategy enables the simultaneous inclusion of multiple DP regions in a single simulation system. In principle, the system potential energy of DP/MM equals the sum of MM and DP as follows:

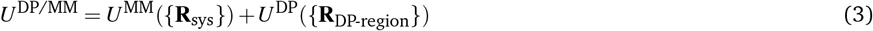

Specifically, the DP region is determined in two steps: (1) selection of the zinc ions and their coordinated atoms based on a distance criterion (3.5 Å is used in this work), and (2) extension from all selected atoms to the corresponding full residues. The coordinates of all atoms in the DP region are fed into the DP model. However, they are not sufficient to construct the full sets of atom environment vectors required to reliably forecast atomic forces without loss of ambient information. This inadequacy arises because the DP regions are often considerably smaller than the entire system, and atoms near the boundaries of the DP region could have atomic environment vectors that lack information about spatially adjacent atoms that fall outside the DP region. To overcome this issue, DP-predicted atomic forces are only applied to the inner atoms in the DP region. Specifically, during the inference of the DP model for zinc-containing proteins in MD simulations, the DP model within the DP/MM framework is effective only for the zinc ions, side chain heavy atoms of residues coordinating zinc, and water oxygen atoms in the DP region, while the DP-predicted atomic forces on the backbone of each coordinated residue are discarded. As a result, the energy prediction provided by the DP model is also ignored during the inference.

### 2.3 Training of DP model for DP/MM

A well-trained MLP depends on three essential factors: (i) a symmetry-preserving architecture that adheres to the principle of physical laws, (ii) efficient sampling of representative chemical structures for preparing the training dataset, and (iii) a scene-oriented loss function that guides the training of MLP towards practical applications. The architecture of DP fulfills (i), being invariant to rotation, translation, and permutation with respect to the input coordinates. Thus, the training of the DP model for DP/MM primarily focuses on the preparation of the training data and the design of the loss function. To extract representative zinc-residue coordinating structures, we systematically searched for zinc-containing proteins in the Protein Data Bank (PDB) database and fed the selected coordination structures into QM calculations to generate training data. Furthermore, the loss function for the DP model was formulated to include the difference in atomic forces between the MM and the QM calculations, ensuring QM-level accuracy with MM-level efficiency for DP/MM simulations.

The experimental structures of zinc-containing proteins in the PDB database serve as the primary source for the training set to cover the chemical space of zinc-coordinated structures in proteins. We retrieved the coordinates of the zinc ions, as well as their coordinated residues and water molecules, from all crystal structures of zinc-containing proteins whose resolution is better than 4 Å. Selection of coordinated atoms was based on a distance criterion of 3.5 Å to the zinc ion, and only coordination groups that include cysteine, histidine, aspartic acid, glutamic acid, water molecules and zinc ions were chosen. In addition to these conformations retrieved from PDB, we supplemented the data set with selected structures picked from simulation trajectories for various zinc-containing proteins and zinc in an aqueous solution.

Using retrieved zinc coordination structures from PDB only, 20,417 conformations were fed into training for a preliminary DP model. However, employing this model in DP/MM simulations led to unsatisfactory results. In most cases, the coordination structure deformed within 1 ns simulation and the zinc ion detached from the protein. To address this issue, non-equilibrated coordination structures were extracted from these preliminary DP/MM simulation trajectories to supplement the training dataset. In particular, structures that captured the ligand exchange process within the coordination group were selected. Once a ligand exchange event was identified, the coordination structures were extracted at the interval of 1 ps. In total 3,983 conformations were generated from these preliminary trajectories, whose coordination groups include *HSD*^3^*Glu, His*^3^*Wat*^3^, *Cys*^3^*HSD, Cys*^4^, and *Cys*^2^*HSD*^2^. In addition, 1,618 zinc-waters coordination structures were generated from MD simulations of a zinc ion solvated in an aqueous solution using the CHARMM force field. 22 sets of 10 ns NVT simulations were performed at temperatures from 303 K to 3400 K, to sample various zinc-water conformations. The number of coordinated water molecules in these structures ranged from 3 to 10. Finally, these three sets were combined to form the final training and validation dataset, comprising a total of 26,018 structures.

The DP model was trained with the difference in atomic forces as calculated by MM and QM methods. MM forces were computed using the CHARMM36m force field ^67^, while B3LYP-D3BJ/cc-pVTZ was adopted for QM calculations. QM calculations were performed with the ORCA package ^68^. Input structures for both MM and QM calculations were preprocessed using CHARMM ^69^ for saturation and relaxation. First, hydrogen atoms and N-methyl groups were patched to the amino and carboxyl groups of selected residues, respectively.

For consecutive residues connected by peptide bonds, only the unsaturated terminals were patched. Coordination structures were then saturated using the *hbuild* command in CHARMM, followed by a geometric relaxation of hydrogen atoms and residue backbone atoms through 500 steps of steepest descent (SD) optimization. These prepared coordination structures were fed into the CHARMM and ORCA for MM and QM calculations, respectively.

To align with the intended application, the loss function used in training the DP model incorporates the QM and MM differences in the forces of zinc ions, water oxygen atoms, and heavy atoms in the side chains of residues. Forces exerted on the hydrogen atoms were excluded from the loss function to avoid bias arising from their inherent fluctuations. Exclusion was implemented via the atom force pre-factor *m*_*i j*_ as outlined in Eq. 4. The variable *n*_*i*_ denotes the number of atoms in the *i*-th structure in the batch, and *m*_*i j*_ represents the value of the pre-factor for the *j*-th atom in the *i*-th structure. It equals 1 if the atomic forces for *j*-th atom were included in the training and 0 if not.

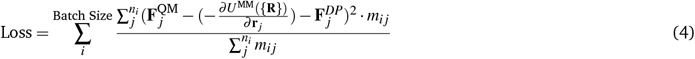

We further note that the original DP architecture was designed to model all the particles in relatively homogeneous condensed phase systems. In contrast, the DP/MM protocol applies the DP model to much smaller regions that dynamically change, containing varying numbers of atoms. For this purpose, we implemented a more concise DP model using self-defined operators that are capable of handling input structures with a varying number of atoms in simulations. In particular, we address the inconsistency in input size by padding the input structures with dummy atoms. To remove the impact of these dummy atoms on the training process, we modify the weighted factor *s*(*r*_*i j*_) for neighbor atom *j* centered on atom *i*, setting it to 0 if either *j* or *i* is a dummy atom. Recall that *s*(*r*_*i j*_) is defined as 1*/ ∥***r**_*i j ∥*_, where **r**_*i j*_ is the relative position vector between atom *i* and *j*, setting *s*(*r*_*i j*_) to 0 effectively positions the dummy atoms infinitely far from the real ones in the DP model. Furthermore, the forces of dummy atoms are ignored during training by setting the corresponding pre-factors to 0. Our extension of this concise DP architecture is detailed in Figure S1 in the Supporting Information and has been integrated into the most recent version of DeePMD-kit ^70^.

### 2.4 DP/MM Simulations

To evaluate the performance of the DP/MM approach in capturing zinc-protein interactions, we carried out simulations on six different zinc-containing proteins as well as on zinc in aqueous solutions (Table S3). The six protein systems feature distinct coordination groups including *Cys*^4^ (6ZFV), *Cys*^3^*His*^1^ (7EEZ), *Cys*^2^*His*^2^(1SP2), *His*^3^*Wat*^3^ (1MSO), *His*^3^*Wat*^1^ (1CA2), and *His*^2^*Glu*^1^*Wat*^1^ (5CPA). All simulations were run by OpenMM ^71^ with a DP/MM plugin we implemented to utilize trained DP models as a new “DeepmdForce” class. The plugin necessitates a DP model and the residue IDs of the zinc ions for initialization. For systems containing multiple spatially separated zinc ions, the DP/MM framework is versatile enough to simultaneously simulate multiple DP-regions.

During the simulations, DP-regions are dynamically updated based on the distance to the zinc ions. In alignment with the composition of the training dataset, only cysteine, histidine, glutamic acid, aspartic acid, and water molecules were considered in the DP-region. There is a small disparity in the input chemical space between the training dataset and the inference processes, which involves the hydrogen and N-methyl groups used to patch terminals during the training. The DP/MM plugin doesn’t implicitly patch such that these atoms remain as backbone atoms in adjacent residues, as we hypothesize that their impact on zinc-protein interactions is negligible. These atoms were only used to construct the environmental descriptor for inner atoms that experience forces computed by the DP model.

All systems were constructed using CHARMM-GUI ^72^, and 200 steps SD minimization were carried out. Atoms in the protein backbone were restrained by harmonic forces with a force constant of 4.0 *kJ/*(*mol ·* Å^2^), while side-chain atoms were restrained with a force constant of 0.4 *kJ/*(*mol ·* Å). For each system, 1 ns NVT simulations were carried out with a 1 fs time step, using either the DP/MM model or the classical C36m FF. This is long enough for comparing the two approaches, while for some systems DP/MM simulations were extended to 10 ns to demonstrate stability. Langevin thermostat was utilized for temperature control at 300 K, and the friction coefficient was set to 1 *ps*^*−*1^. Detailed information on the simulation systems is provided in Table S3.

## 3 Results

### 3.1 Dataset Component Analysis and Training Result

The dataset contains a total of 26,018 zinc-coordination conformations, with the majority (20,417) sourced from protein crystal structures. The remaining 5,601 conformations were derived from preliminary simulations, 3,983 from zinc-containing proteins and 1,618 from zinc-water systems. Within these 26,018 conformations, cysteine appeared 49,727 times, accounting for 48.4% of the selected residues. Histidine constituted the second most common category (24.3%). Significant variation between the two protonated forms of histidine was observed: 17.1% were *δ* -nitrogen protonated (HSD), and 7.2% were *ε*-nitrogen protonated (HSE). Water molecules comprised 18.8% of the selected residues, while the remaining residues were predominantly glutamic acid (3.8%) and aspartic acid (4.1%), as illustrated in Fig. 1a. In terms of coordination groups (Fig. 1b), cysteine-rich categories were prevalent. Specifically, the *Cys*^4^ group constituted 24.4% (6,343) of all conformations. The *Cys*^3^*HSE*^1^, *Cys*^3^*HSD*^1^ and *Cys*^2^*HSD*^2^ groups accounted for 9.5%, 6.1% and 4.1%, respectively, leaving the remaining 55.9% distributed among 350 other coordination groups. The addition of simulation-derived conformations enriches the representation of these less prevalent coordination groups, as shown in Fig. S2.

**Fig. 1.**
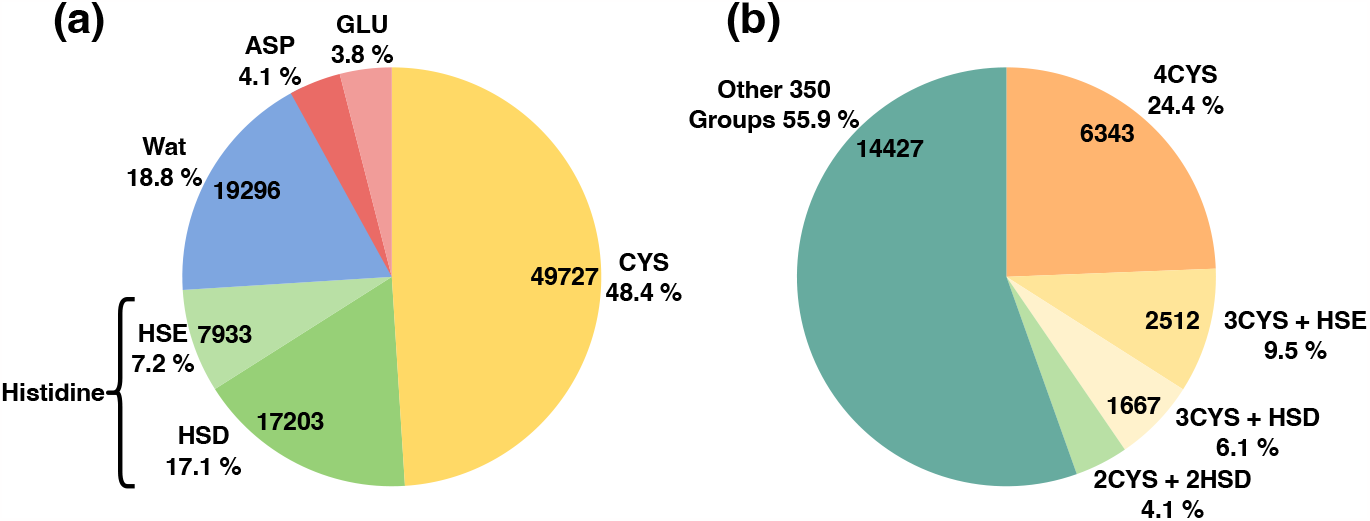
Pie chart displaying the occupancy of selected residues (a) and coordination groups (b) in the final dataset for training and validation.

The dataset was randomly split into training and test sets, containing 25,954 and 64 structures, respectively. The training set was further divided with a 9:1 ratio to be used for training and validation. The training process was capped at 1.5 million steps with an initial learning rate of 0.001, which decayed by a factor of 0.98 every 20,000 steps. The training loss function, defined as the sum of force differences over the atoms of interest, is detailed in Section 2.3. Upon completion of 1.5 million training epochs, the training loss (RMSE) reached 2.5 kcal/(mol *·* Å), and the test loss registered at 3.0 kcal/(mol *·* Å). Utilizing this trained DP model, subsequent DP/MM simulations outperformed their MM counterparts in modeling the structures of zinc-containing proteins, as presented below.

### 3.2 DP/MM Maintaining the Preferred Tetrahedral *Cys*^*x*^*His*^*y*^ Coordination Groups

We first elucidate the accuracy of DP/MM simulations for major coordination groups, denoted as *Cys*^*x*^*His*^*y*^ (where *x* + *y* = 4), in which the zinc ion establishes tetrahedral coordination structures with deprotonated sulfur atoms of cysteine residues and nitrogen atoms of histidine residues. The N-terminal zinc finger domain of the GATA2 transcription factor (PDB ID: 6ZFV) serves as an example featuring one zinc coordinating with a *Cys*^4^ group. Additional examples, including the SHH2 SAWADEE domain (PDB ID: 7EEZ) and the SP1F2 zinc finger domain (PDB ID: 1SP2), are employed to assess zinc interactions with the *Cys*^3^*HSD* and the *Cys*^2^*HSD*^2^ coordination groups, respectively.

A 1 ns MM simulation of 6ZFV using the C36m FF demonstrated that a water molecule intruded upon the coordination center, shifting the coordination structure from *Cys*^4^ to a *Cys*^4^*Wat*^1^ group, forming a bipyramidal geometry. It is well known that inaccurate modeling of zinc interactions in classical force fields can lead to unphysical over-coordination by water. The increase in CN from 4 to 5 and the alteration of coordination geometry occurred within tens of ps, as depicted in Fig. 2a. Concurrent with water binding, the distance between the zinc ion and the S*γ* atom of Cys48 increased by 0.2 Å, deviating further from the value observed in the crystal structure (Fig. 2d).

**Fig. 2.**
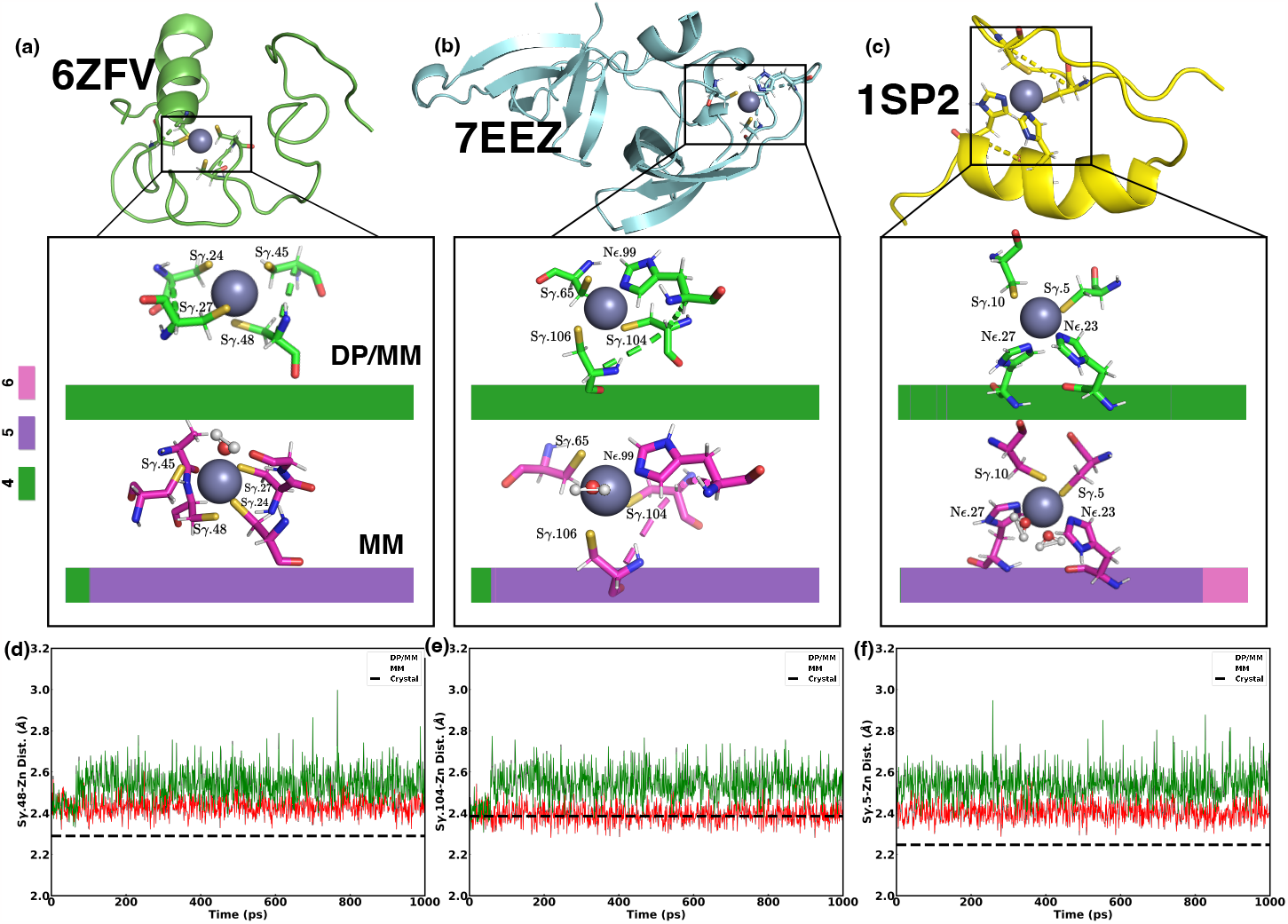
The coordination structures of the zinc ion in proteins 6ZFV (a), 7EEZ (b) and 1SP2 (c), plotted using the last frame from 1 ns DP/MM and MM simulations. The zinc ion is represented as a silver-gray ball. Accompanying color bars elucidate the CN along the 1 ns simulations. These CN values quantify the number of atoms directly coordinating with the zinc ion: green indicates a CN value of 4, purple signifies 5, and pink denotes 6. In addition, the time evolution of representative distances between the zinc ion and its directly coordinated atoms is presented: (d) S*γ*.48-Zn in 6ZFV;(e) S*γ*.104-Zn in 7EEZ; (f) S*γ*.5-Zn in 1SP2.

In contrast to conventional MM simulations, the DP/MM simulation of 6ZFV yielded a compact tetrahedral coordination geometry resembling the *Cys*^4^ group, as evidenced in Fig. 2a. More accurate and balanced interactions based on the DP/MM approach effectively eliminated the problem of excessive water binding. The distance of the nearest water molecule relative to the zinc ion fluctuated between 4 and 6 Å (Figure S3d). Consequently, the distances between zinc and its directly coordinated sulfur atoms were shorter with DP/MM and aligned more closely with crystal measurements (see Table 1 and Figure 2c and S3). The fluctuation of these interatomic Zn-S*γ* distances was also slightly reduced. To further validate the stability of the DP/MM approach, the simulation was extended to 10 ns, during which the tetrahedral coordination geometry remained intact and the Zn-S*γ* distances remained in agreement with the 1 ns simulation results (Table S1).

**Table 1.** Distances between the zinc ions and their directly coordinated atoms averaged over MD trajectories. The fluctuation of distances computed as standard deviation is shown in parentheses. For the 1CA2 protein, we also include oxygen-oxygen distances within the hydrogen bonding network for comparison. Values that match the crystal values better are highlighted in bold.

| Protein | Dist. Type | MM | DP/MM | Crystal |
| --- | --- | --- | --- | --- |
| 6ZfV | Sγ.24-Zn | 2.452 (0.051) | **2.415** (0.047) | 2.320 |
|  | Sγ.27-Zn | 2.525 (0.072) | **2.464** (0.061) | 2.270 |
|  | Sγ.45-Zn | 2.521 (0.069) | **2.504** (0.066) | 2.318 |
|  | Sγ.48-Zn | 2.536 (0.075) | **2.436** (0.046) | 2.290 |
| 7EEZ | Sγ.65-Zn | 2.466 (0.057) | **2.401** (0.044) | 2.353 |
|  | Sγ.104-Zn | 2.539 (0.075) | **2.415** (0.048) | 2.386 |
|  | Sγ.106-Zn | 2.502 (0.065) | **2.394** (0.043) | 2.232 |
|  | Nε.99-Zn | 2.179 (0.058) | **2.181** (0.065) | 2.207 |
| 1SP2 | Sγ.5-Zn | 2.538 (0.069) | **2.407** (0.046) | 2.247 |
|  | Sγ.10-Zn | 2.445 (0.056) | **2.358** (0.037) | 2.246 |
|  | Nε.23-Zn | **2.170** (0.063) | 2.237 (0.101) | 2.033 |
|  | Nε.27-Zn | **2.165** (0.058) | 2.202 (0.097) | 2.124 |
| 1MSO | Nε-Zn.103 | **2.192** (0.058) | 2.203 (0.075) | 2.093 |
|  | OH2-Zn.103 | 2.111 (0.058) | **2.188** (0.086) | 2.204 |
|  | Nε-Zn.104 | **2.191** (0.058) | 2.203 (0.076) | 2.102 |
|  | OH2-Zn.104 | 2.110 (0.058) | **2.191** (0.088) | 2.234 |
| 1CA2 | Nε.91-Zn | 2.313 (0.096) | **2.136** (0.053) | 1.995 |
|  | Nε.93-Zn | **2.153** (0.049) | 2.155 (0.060) | 2.103 |
|  | Nδ.116-Zn | 2.224 (0.065) | **2.210** (0.072) | 1.910 |
|  | Zn-OH2 | 2.101 (0.057) | **2.143** (0.083) | 2.087 |
|  | OH2-Oγ.195 | 2.903 (0.178) | **2.700** (0.170) | 2.780 |
|  | Oγ.195-Oε1.103 | 2.789 (0.122) | **2.617** (0.111) | 2.433 |
|  | Oγ.195-Oε2.103 | 2.748 (0.138) | **3.613** (0.368) | 3.459 |
|  | Oε1.103-Zn | 2.020 (0.054) | **5.296** (0.506) | 4.060 |
|  | Oε2.103-Zn | 3.874 (0.126) | **5.654** (0.420) | 5.459 |
| 5CPA | Nδ.69-Zn | 2.240 (0.070) | **2.146** (0.056) | 2.126 |
|  | Nδ.196-Zn | 2.224 (0.065) | **2.169** (0.065) | 1.910 |
|  | Oε1.72-Zn | 2.067 (0.068) | **2.045** (0.068) | 2.176 |
|  | Oε2.72-Zn | **2.119** (0.092) | 2.009 (0.052) | 2.307 |
|  | OH2-Zn | **2.109** (0.060) | 2.165 (0.083) | 2.049 |

Similar observations were obtained when comparing MM and DP/MM simulations of the 7EEZ protein. With the MM force field model, one water molecule was attracted and bound to the zinc ion after about 50 ps of simulation, distorting the initial *Cys*^3^*HSD* tetrahedral group and forming a bipyramidal coordination geometry (Fig. 2b). Meanwhile, the DP/MM simulation for 7EEZ preserved the tetrahedral *Cys*^3^*HSD* coordination group along 1 ns or 10 ns (Table S1) simulations. Similar to the 6ZFV results, the Zn-S*γ* distances in the DP/MM simulation were shorter and compared favorably with the crystal structure values, as shown in Fig. 2e and Table 1. In contrast, the average distances between the zinc ion and the N*ε* atom of HIS99 were the same between MM (2.179 Å) and DP/MM (2.181 Å), both close to the crystal value (2.207 Å). This implies that the interaction between zinc and the coordinated histidine might be stronger and less susceptible to the influence of excessive water binding.

The zinc finger domain 1SP2 contains one zinc coordinated with two cysteine and two histidine residues. In the MM simulation of 1SP2, water molecules were more likely to be attracted to the coordination sphere. The CN value immediately rose to 5 and, after about 800 ps, further increased to 6, as two additional water molecules coordinated with the zinc ion by the end of the 1 ns simulation (Figure 2c). With DP/MM, the tetrahedral coordination involving the *Cys*^2^*HSD*^2^ group remained stable during 1 ns MD simulation. In several instances, water can approach and coordinate with zinc in a transient manner. For example, at 139 ps a water molecule came within 2.5 Å of the zinc ion, but unlike in MM simulations, the water left after 2 ps (Figure S5d). This indicates that water can access the coordination center for the *Cys*^2^*HSD*^2^ group but does not remain bound, allowing the tetrahedral coordination geometry to be preserved using the DP/MM approach.

MD simulations of 6ZFV, 7EEZ and 1SP2 proteins demonstrated the robustness of the DP/MM approach in accurately modeling zinc-CYS/HSD dynamics. The Zn-S distances in DP/MM simulations are shorter and notably closer to crystal structure values compared to those obtained using conventional MM models. The Zn-N*ε* distances with DP/MM are similar or slightly larger than the MM values. In terms of coordination geometry, DP/MM consistently produced tetrahedral coordination with the *Cys*^4^, *Cys*^3^*HSD*, and *Cys*^2^*HSD*^2^ groups, contrasting starkly with the over-hydrated geometries that result from MM simulations a phenomenon previously linked to inaccuracies in classical FFs ^23,28^. The over-binding of water was previously attributed to the inaccuracy of zinc-sulfur interactions due to the absence of explicit charge transfer in classical force fields ^21,28,43^. Recent work has shown that machine learning potentials might effectively capture charge transfer effects ^59^, corroborating our observations of accurate tetrahedral coordination in MD simulations adopting the DP/MM model.

### 3.3 Validation on *His*^3^*Wat*^*x*^(*x* = 3 or 1) Coordination Groups

The T6 human insulin (PDB ID: 1MSO) presents a particularly interesting case as it features two zinc ions, each octahedrally coordinated with the *HSD*^3^*Wat*^3^ group. In this study, the two zinc ions were placed in two separate DP regions and simultaneously modeled in the DP/MM simulation. 1 ns MD simulations using either the MM or the DP/MM models successfully replicated the octahedral coordination geometry with the *HSD*^3^*Wat*^3^ group, aligning well with the crystal structure. Comparing the distances between zinc and its coordinated atoms, we note that for both zinc ions, the three histidine N*ε*-Zn distances and the three water O-Zn distances are identical in the crystal structure. This is likely due to manual constraints applied during model building. Nevertheless, our simulation results indicated that these distances were indeed very similar (Table S2) so the averaged values are compared in Table 1. In alignment with the *Cys*^*x*^*His*^*y*^ results, the distances between zinc and N*ε* atoms were almost the same between MM and DP/MM, with the DP/MM distances being slightly larger. Both zinc ions are coordinated with three water molecules, and it was observed that these coordinating water molecules remained stably bound in the DP/MM simulation. Their distances relative to zinc were 0.08 Å larger than those obtained in the MM simulation, bringing them closer to the crystal structure values.

Zinc can also coordinate with three histidine residues and one water molecule, as exemplified in this study by the carbonic anhydrase II (PDB ID: 1CA2) ^73^. Specifically, two of the coordinating histidine contribute N*ε* atoms, while the third histidine offers an N*δ* atom. Therefore, the coordination group is *HSD*^2^*HSE*^1^*Wat*^1^ considering the protonation states of ligands (Figure 3a). Interestingly, the crystal structure reveals that the coordinating water molecule is stabilized by a hydrogen bonding network (Wat258-Thr195-Glu103), which has been demonstrated to be essential for the protein’s enzymatic activity ^24,73–75^.

**Fig. 3.**
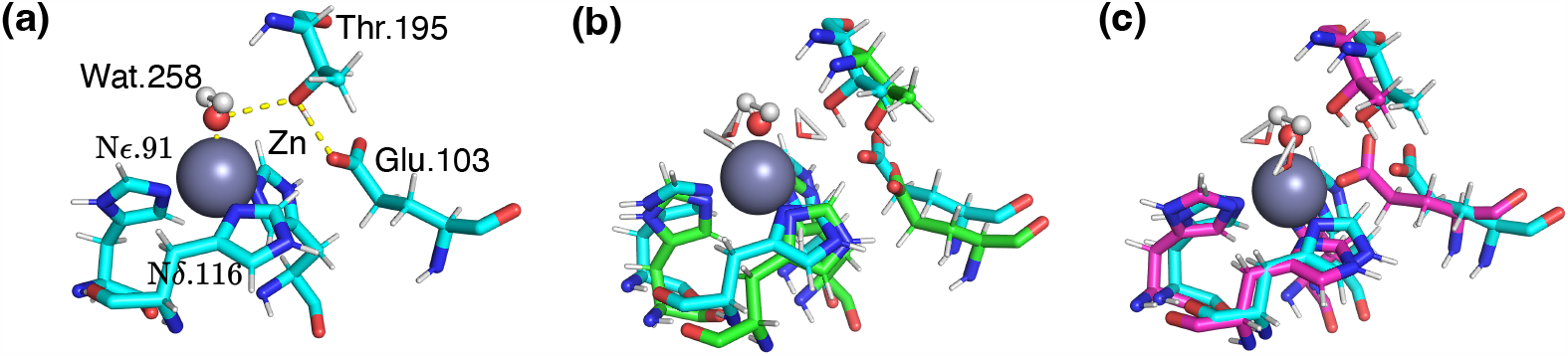
The coordination geometry of 1CA2. (a) The crystal structure, with the hydrogen bonding network of Wat258-Thr195-Glu103 highlighted by yellow dashed lines. (b) The structure extracted from the last frame of 1 ns DP/MM trajectory (green) aligned with the crystal structure (cyan). (c) The structure extracted from the last frame of 1 ns DP/MM trajectory (magenta) aligned with the crystal structure (cyan). The coordinating water molecule in the crystal structure is depicted using a ball-and-stick representation, while the coordinating water molecules in simulations are represented by licorice.

During the MM simulation of 1CA2, this crucial hydrogen bonding network was disrupted by the attraction between the zinc ion and the glutamic acid (Glu103). At the very beginning of the simulation, Glu103 was drawn into the coordination sphere. Along with two water molecules and three histidines, the O*ε*1 atom of the carboxyl group in glutamic acid formed an octahedral coordination group around the zinc ion (Figure 3c). In contrast, during the DP/MM simulation the carboxyl group did not engage in the coordination of zinc, and the hydrogen bonds between one coordinating water and Thr195, as well as between Thr195 and Glu103, were well preserved (Figure 3b). As a consequence, the coordination geometry was better maintained.

It’s interesting to note that 1CA2 is one of the two zinc-containing protein systems used to test the CHARMM non-bonded model for zinc by Stote and Karplus ^24^. Our observation that an improper coordination structure disrupted the hydrogen bond network with MM is consistent with their findings. Regarding coordination numbers, both MM and DP/MM simulations resulted in a CN of 6, larger than the CN of 4 observed in the crystal structure (Figure S7). This discrepancy may be attributed to differences in protonation states ^76^. Despite the additional coordination of water molecules, the hydrogen bond network atom pairs in the 1CA2 crystal structure were well preserved in the DP/MM simulation, and the distances between these pairs were closer to those in the crystal structure, as illustrated in Table 1.

### 3.4 Validation on the Interaction between Zinc and Carboxyl Groups

The preservation of the key hydrogen bonding network in the DP/MM simulation of 1CA2 is encouraging and indicates that the DP/MM model does not overestimate the interaction between zinc and carboxyl side chain groups. To further validate these types of interactions, we carried out MD simulations of carboxypeptidase A (PDB ID: 5CPA) ^77^, another zinc-containing protein previously investigated by Stote and Karplus ^24^. In 5CPA, a single zinc ion is coordinated with the carboxyl group from one glutamic acid, a water molecule, and two N*δ* atoms from two histidine residues, constituting a *HSE*^2^*Glu*^1^*Wat*^1^ group. We note that both oxygen atoms of the carboxylate ligand are engaged with the zinc ion together in 5CPA such that it’s actually in a bidentate tetrahedral geometry (Figure 4a). As a consequence, we count the coordination number to be 5 in the crystal structure, rather than the typical 4 in a standard tetrahedral coordination, given that the glutamic acid contributes two liganding atoms.

**Fig. 4.**
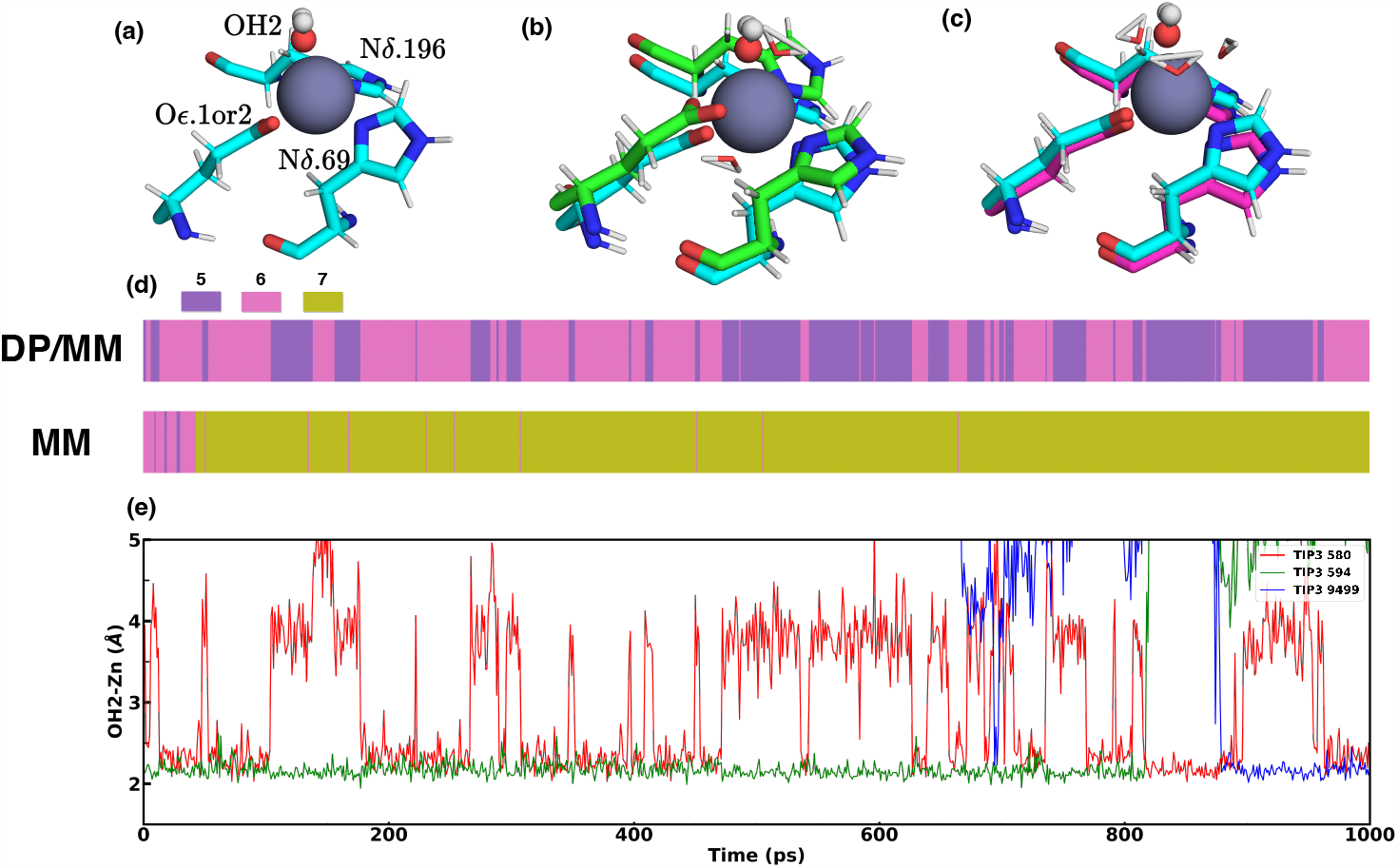
The coordination geometry of 5PCA in the crystal structure (a) and the last frame of 1 ns DP/MM (b) and MM (c) trajectory. The crystal structure is colored in cyan, while the DP/MM and MM structures were aligned to the crystal structure and colored in green and magenta, respectively. The coordinating water molecule in the crystal structure is depicted with ball-and-stick, while those in simulations with licorice. The variation in the number of coordinated water molecules between DP/MM and MM is also depicted in the time evolution bar of CN in panel (d). We note that since both oxygen atoms of the carboxyl ligand are coordinated with the zinc ion in the crystal structure, the CN value for the bidentate tetrahedral coordination group in the crystal structure is considered to be 5 (colored in purple). Correspondingly, a CN value of 6 (pink) or 7 (olive) indicates that there are two or three water molecules being coordinated. To highlight the dynamics of the coordinated water molecules in the DP/MM simulation of 5CPA, the distances between these water molecules and the zinc ion are presented in panel (e).

During the MM simulation of 5CPA, the CN values first increased to 6 and then to 7, with three water molecules observed coordinated with the zinc as shown in Fig. 4c. These water molecules displayed dynamic behavior, spinning around the zinc ion and frequently exchanging positions with nearby water molecules, a phenomenon highlighted by the strip pattern in the time evolution bar of CN values (Fig. 4d). In the final frame of the DP/MM simulation, two water molecules were found to be coordinated with the zinc ion, as illustrated in Fig. 4b. These molecules were positioned on opposite sides of the zinc ion, causing a relatively large conformational change in the side chains of the coordinated residues. It’s worth noting, however, that these two water molecules did not maintain stable coordination throughout the 1ns DP/MM simulation. The number of coordinated water molecules alternated between one and two (Fig. 4d). For half the duration of the DP/MM simulation, a bidentate tetrahedral coordination group, similar to that observed in the crystal structure, was formed with one water molecule.

The exchange of water molecules during the DP/MM simulation is detailed in Fig. 4e. One water molecule (wat594) remained coordinated at a position similar to that of the crystal water for 800 ps, while another water molecule (wat580) alternated between bound and unbound states at the opposite position. When wat594 dissociated from the coordination sphere, wat580 can bind stably. In the meantime, a water molecule initially present in the solution (wat9499) approached and subsequently occupied the coordination position of the crystal water, thereby replacing the original coordinated water molecule. Geometric measurements of these coordination structures in MM and DP/MM simulations are compared in Table 1. The N*δ* -Zn distances in the DP/MM simulation were shorter than those in the MM simulation and closer to the crystal values. We note that the carboxyl group from glutamic acid and aspartic acid were slightly underrepresented in our training data set, collectively accounting for 7.9 % of the coordinated residues selected. Nevertheless, MD simulations of 5CPA indicated that the interactions between zinc and carboxyl groups were accurately described by the DP/MM model. They also demonstrated that the ligand exchange process can occur with molecules entering and leaving the DP region during the simulation.

### 3.5 Performance on Solvated Zinc Ions

In the simulations of zinc-containing proteins previously described, we demonstrated that the DP/MM approach effectively alleviates the over-coordination of water molecules in the zinc coordination structure. We assume that this is due to the more accurate manybody effects that balance the interactions between the zinc ion, amino acids, and water molecules, rather than simply weakening the zinc-water interactions. To further illustrate the impact of DP/MM on zinc-water interactions, we carried out MD simulations using an aqueous-solvated zinc ion system. The system comprises a single zinc ion solvated in a cubic water box containing 572 water molecules. The radial distribution function (RDF), or g(r), of water oxygen atoms relative to the zinc ion was computed from 20 ns NVT simulations at 300 K. As depicted in Fig. 5a, the RDFs obtained with the DP/MM and the MM models closely resemble each other. Specifically, the first peaks occur at 2.138 Å and 2.106 Å for DP/MM and MM, respectively. Notably, the first peak in the DP/MM RDF is closer to both the experimental value of 2.14 Å ^78^ and the *ab initio* molecular dynamics results (2.18/2.19 Å) ^79^. This slight shift towards a larger peak in the DP/MM simulation suggests a subtle weakening of the zinc-water interaction. The oxygen-oxygen distances in the first hydration shell were also quite similar in both models, measuring 2.972 Å in each case. Furthermore, the running integration of the RDFs at 300 K revealed that both DP/MM and MM models indicate six coordinated water molecules within the first hydration shell of the zinc ion, which aligns with experimental findings (Figure 5a). These results indicate that the DP/MM approach accurately captures the static structural arrangement of water molecules around the zinc ion.

**Fig. 5.**
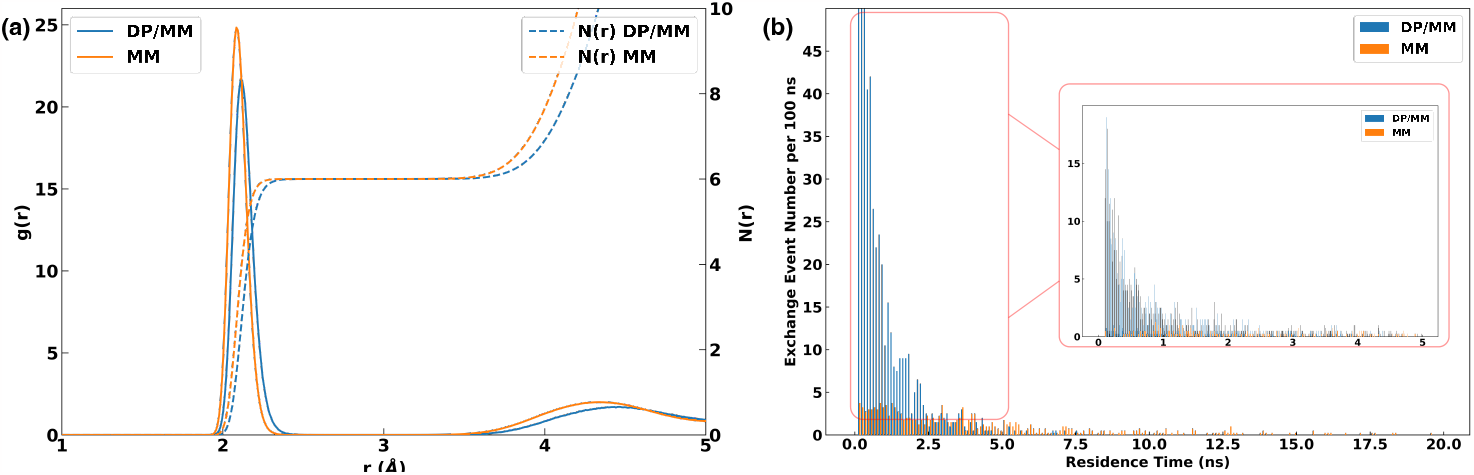
(a) Radial distribution function, or g(r), of water oxygen atoms relative to the zinc ion computed using DP/MM and MM simulations at 300 K. N(r), which represents the number of coordinated water molecules within distance *r* from the zinc ion, is also plotted as the running integration of g(r). (b) Histogram distribution of residence time for water molecules in the first solvation shell of zinc at 350 K with a bin size of 0.1 ns. The inset in (b) displays the histogram of water residence times ranging from 0 to 5 ns with a bin size of 0.01 ns.

We also characterized the dynamic impact of zinc-water interactions by estimating the residence time of water molecules in the first hydration shell of zinc. To accumulate enough water exchange events for statistical analysis on the duration of water molecules in direct contact with zinc, simulations were conducted at an elevated temperature of 350 K to increase the likelihood of water exchange. Two separate 100 ns DP/MM simulations were carried out to estimate the residence time of water molecules, while with MM the trajectories were extended to 200 ns due to the slower rate of water exchange, as discussed below. Water molecules located within 2.5 Å of the zinc ion were considered part of its hydration shell, and their lifetime were recorded with a buffer window of 100 ps. Specifically, an exchange event was only counted if a water molecule left the first hydration shell and remained outside of it for at least 100 ps.

As illustrated in Fig. 5b, DP/MM simulations yielded a higher frequency of water exchange events compared to those using MM. Moreover, the residence time of water molecules within the first solvation shell of zinc was shorter in DP/MM. The average residence time was estimated to be 0.8 ns and 4.2 ns with the DP/MM and MM trajectories, respectively. The reduced residence time in DP/MM simulations indicates more dynamic behavior for water molecules in direct contact with zinc. It’s also important to note that both DP/MM and MM simulations reported residence times on the same ns timescale, suggesting that the improvement brought by DP/MM does not fundamentally alter the zinc-water interaction dynamics.

## 4 Conclusions and Discussion

In this study, we present a unique approach that integrates machine learning potential with classical force fields. The classical FF, or the MM model, is used to describe interactions in the entire system, while an MLP is used to augment the predicted forces on atoms within an adaptively selected region of interest. We choose zinc-containing proteins as test cases to demonstrate the effectiveness of this approach, as these are known to present challenges for classical FFs in accurately describing coordination numbers and geometries. More specifically, we implemented our method in OpenMM and referred to it as DP/MM, which uses the Deep Potential model as the MLP to bridge the gap between the classical FF (CHARMM36m) and QM calculations (B3LYP-D3BJ/cc-pVTZ). Validation on a variety of zinc-containing protein systems confirms that our proposed method can accurately reproduce the coordination geometry of zinc ions. Our results demonstrate the power of integrating MLPs with FFs to model and simulate metalloprotein systems. The DP/MM approach can be readily extended to simulate other metalloprotein systems, provided that appropriate training sets are generated. It might also be utilized to enhance the accuracy of modeling ligand-protein interactions.

Six zinc-containing protein systems, each featuring a specific coordination group, along with an aqueous zinc system, were employed to validate the performance of DP/MM. Notably, DP/MM succeeded in accurately modeling the coordination geometries along MD simulations, outperforming MM in these test systems. In the simulations of *Cys*^4^, *Cys*^3^*HSD* and *Cys*^2^*HSD*^2^ coordination groups of zinc ions, DP/MM consistently generated the correct tetrahedral coordination geometry. This stands in contrast to MM simulations, which were unable to establish this geometry due to the over-coordination of water molecules. Furthermore, DP/MM exhibited balanced modeling of zinc-water interactions in several systems whose coordination groups included water molecules. This is evidenced by the OH2-Zn distances that are closely aligned with crystallographic data and the observation of bidentate tetrahedral coordination groups involving water. When compared to MM simulations for the 1CA2 protein, DP/MM preserved a functionally crucial hydrogen bond network that held one water molecule in coordination with zinc. These results suggest that more reasonable structures and dynamics can be obtained by modeling the atomic forces in the region of interest at the QM level via the DP/MM approach.

In addition to achieving high accuracy in simulating zinc-protein interactions, DP/MM offers several other noteworthy advantages. The first of these is a remarkable reduction in the computational scaling and costs traditionally associated with QM calculations, effectively bringing them down to the MM level. Leveraging the computational efficiency of the DP model, simulations of typical solvated protein systems can run at the rates of several ns per day. For instance, using one NVIDIA GTX1080Ti GPU card along with 4 CPU cores, the DP/MM simulation for the 6ZFV protein (18254 atoms in total) runs at 1.71 ns/day. In scenarios involving multiple DP regions in a single DP/MM simulation, such as the 1MSO protein with two zinc-containing DP regions, a rate of 0.61 ns/day was achieved using the same GTX1080Ti card. It’s important to note that while the major computational cost lies in machine learning model inference, ample opportunities remain for optimizing the speed of DP/MM simulations. These include minimizing the communication overhead between the MD engine and the MLP calculations, as well as leveraging the engine’s neighbor list array for adaptively selecting the DP region, instead of maintaining its own list.

Second, DP/MM represents a flexible approach that can be readily extended to various applications. This flexibility manifests in two aspects. The first aspect is the adaptability of the model to dynamically changing DP regions, allowing atoms to enter or leave these regions. The ΔE-MLP/MM methods employ MLPs to elevate the quality of the QM region from semi-empirical to *ab initio* QM levels ^63,80,81^. This strategy can be viewed as a hybrid MLP+QM/MM scheme, albeit one that remains reliant on and constrained by existing QM/MM methods. The second aspect is that the model obviates the need for excluding bonded or non-bonded MM interactions within the DP region, a requirement in many existing MLP/MM frameworks. Previously proposed MLP/MM often necessitate elaborate exclusion rules for bonded and/or non-bonded MM interactions when atoms are described by MLPs ^82–84^. In contrast, our DP/MM model uses DP to compute supplementary corrections to the MM atomic forces. We demonstrate that there is no specific need for interaction exclusions within the DP regions as the intramolecular and intermolecular interactions from FFs are integrated during the training of the DP model. This further simplifies the integration of DP/MM with standard MD engines.

The third advantage of DP/MM demonstrated in this work is its generalization ability. A unified DP model can handle a variety of different zinc coordination groups, thanks to the extensive parameter space inherent in neural networks. We provide this well-trained model as a new type of FF modification. Looking ahead, publicly available force fields may transition from textual bonded and non-bonded parameters to trained MLPs ^85^. It is worth noting that the transferability of the DP model is primarily constrained by the diversity of the training dataset. Therefore, the integration of more robust MLP training schemes, such as active learning ^86^ or the use of pre-trained large models, could further enhance DP/MM’s ability to generalize to more complex coordination situations.

Despite the many advantages of the DP/MM scheme, its limitations should also be discussed. As only the interior particles of the DP region are subject to the added atomic forces, there is no trivial way to obtain the potential energy from the DP model. Consequently, we are not able to perform free energy calculations or examine the thermodynamic properties using DP/MM. To obtain valid energy estimations that are consistent with the added atomic forces in DP/MM, future work will require rigorous implementation of numerical integration on these forces, along with comprehensive testing.

A scenario that the current DP/MM scheme cannot model involves proteins containing a cluster of closely placed zinc ions ^87^. While the DP framework can in principle handle multiple zinc ions within a single DP region, constructing the corresponding training dataset is significantly more demanding. Similarly, the current model is not able to treat small ligands and metabolites coordinated with zinc, due to their absence in the training dataset. Another limitation is the lack of long-range polarization effects in the current DP/MM scheme. To address this issue, one might need to go beyond additive FFs for the MM component; however, natural integration of the DP model and a non-additive FF model remains a challenging task. Despite these limitations, this study demonstrates that DP/MM can accurately model zinc-containing proteins in a versatile and computationally efficient manner. The DP/MM approach presented here provides a promising pathway for extending its applicability to other metalloproteins or enzymatic systems that demand QM-level accuracy.

## Data Availability Statements

The code for DP/MM simulations is available at https://github.com/JingHuangLab/openmm_deepmd_plugin. An example DP/MM simulation script for the 6ZFV protein is also provided in this repository.

## Author Contributions

J.H. conceived the study. Y.D performed all the coding and calculations. Y.D and J.H wrote the manuscript together.

## Acknowledgements

We thank Dr. Yang Guo for his valuable advice. The work is supported by the “Pioneer” and “Leading Goose” R&D Program of Zhejiang (2023C03109) and the National Natural Science Foundation of China (32171247, 21803057). We thank the Westlake University Supercomputer Center for computational resources and related assistance.

## Notes and references

1 J. A. Tainer, V. A. Roberts and E. D. Getzoff, Current Opinion in Biotechnology, 1992, 3, 378–387.

2 T. Dudev and C. Lim, Chemical reviews, 2014, 114, 538–556.

3 M. M. Yamashita, L. Wesson, G. Eisenman and D. Eisenberg, Proceedings of the National Academy of Sciences, 1990, 87, 5648–5652.

4 J. A. Tainer, V. A. Roberts and E. D. Getzoff, Current Opinion in Biotechnology, 1991, 2, 582–591.

5 J. Anastassopoulou and T. Theophanides, Bioinorganic chemistry, Springer, 1995, pp. 209–218.

6 C. Andreini, I. Bertini, G. Cavallaro, G. L. Holliday and J. M. Thornton, JBIC Journal of Biological Inorganic Chemistry, 2008, 13, 1205–1218.

7 C. Guo, M. Cheng and M. L. Gross, Analytical chemistry, 2018, 91, 1416–1423.

8 A. Khandelwal, V. Lukacova, D. Comez, D. M. Kroll, S. Raha and S. Balaz, Journal of medicinal chemistry, 2005, 48, 5437–5447.

9 C. Andreini, L. Banci, I. Bertini and A. Rosato, Journal of proteome research, 2006, 5, 196–201.

10 J. E. Coleman, Annual review of biochemistry, 1992, 61, 897–946.

11 C. Andreini, L. Banci, I. Bertini and A. Rosato, Journal of proteome research, 2006, 5, 3173–3178.

12 W. M. Gommans, H. J. Haisma and M. G. Rots, Journal of molecular biology, 2005, 354, 507–519.

13 H.-W. Sun and B. V. Plapp, Journal of Molecular Evolution, 1992, 34, 522–535.

14 N. M. Hooper, FEBS letters, 1994, 354, 1–6.

15 B. L. Vallee and H. Neurath, J Biol Chem, 1955, 217, 253–261.

16 K. Lindorff-Larsen, S. Piana, R. O. Dror and D. E. Shaw, Science, 2011, 334, 517–520.

17 M. Karplus and J. A. McCammon, Nature structural biology, 2002, 9, 646–652.

18 Y.-P. Pang, Molecular modeling annual, 1999, 5, 196–202.

19 S. Bottaro and K. Lindorff-Larsen, Science, 2018, 361, 355–360.

20 J. Huang and A. D. MacKerell, Curr. Op. Struct. Biol., 2018, 48, 40 –48.

21 R. R. Roe and Y.-P. Pang, Molecular modeling annual, 1999, 5, 134–140.

22 M. Laitaoja, J. Valjakka and J. Jänis, Inorganic chemistry, 2013, 52, 10983–10991.

23 E. Ahlstrand, K. Hermansson and R. Friedman, The Journal of Physical Chemistry A, 2017, 121, 2643–2654.

24 R. H. Stote and M. Karplus, Proteins: Structure, Function, and Bioinformatics, 1995, 23, 12–31.

25 M. Macchiagodena, M. Pagliai, C. Andreini, A. Rosato and P. Procacci, Journal of Chemical Information and Modeling, 2019, 59, 3803–3816.

26 M. Macchiagodena, M. Pagliai, C. Andreini, A. Rosato and P. Procacci, ACS omega, 2020, 5, 15301–15310.

27 L. Banci, Current opinion in chemical biology, 2003, 7, 143–149.

28 N. Calimet and T. Simonson, Journal of Molecular Graphics and Modelling, 2006, 24, 404–411.

29 O. A. Donini and P. A. Kollman, Journal of medicinal chemistry, 2000, 43, 4180–4188.

30 C. R. Landis, T. Cleveland and T. K. Firman, Journal of the American Chemical Society, 1998, 120, 2641–2649.

31 J. Y. Xiang and J. W. Ponder, Journal of computational chemistry, 2013, 34, 739–749.

32 I. Tubert-Brohman, M. Schmid and M. Meuwly, Journal of Chemical Theory and Computation, 2009, 5, 530–539.

33 P. Li and K. M. Merz Jr, Journal of chemical theory and computation, 2014, 10, 289–297.

34 Z. Li, L. F. Song, G. Sharma, B. Koca Findik and K. M. Merz Jr, Journal of Chemical Theory and Computation, 2022, 19, 619–625.

35 R. Wu, Z. Lu, Z. Cao and Y. Zhang, Journal of chemical theory and computation, 2011, 7 2, 433–443.

36 Y.-P. Pang, Proteins: Structure, Function, and Bioinformatics, 2001, 45, 183–189.

37 J. Åqvist and A. Warshel, Journal of molecular biology, 1992, 224, 7–14.

38 F. Duarte, P. Bauer, A. Barrozo, B. A. Amrein, M. Purg, J. Åqvist and S. C. L. Kamerlin, The Journal of Physical Chemistry B, 2014,118, 4351–4362.

39 J. C. Wu, J.-P. Piquemal, R. Chaudret, P. Reinhardt and P. Ren, Journal of chemical theory and computation, 2010, 6, 2059–2070.

40 J. Zhang, W. Yang, J.-P. Piquemal and P. Ren, Journal of chemical theory and computation, 2012, 8, 1314–1324.

41 N. Gresh, J.-P. Piquemal and M. Krauss, Journal of computational chemistry, 2005, 26, 1113–1130.

42 H. Yu, T. W. Whitfield, E. Harder, G. Lamoureux, I. Vorobyov, V. M. Anisimov, A. D. MacKerell Jr and B. Roux, Journal of chemical theory and computation, 2010, 6, 774–786.

43 W. Li, J. Zhang, J. Wang and W. Wang, Journal of the American Chemical Society, 2008, 130, 892–900.

44 D. V. Sakharov and C. Lim, Journal of the American Chemical Society, 2005, 127, 4921–4929.

45 C. E. Tzeliou, M. A. Mermigki and D. Tzeli, Molecules, 2022, 27, 2660.

46 U. Ryde, Current opinion in chemical biology, 2003, 7, 136–142.

47 W. Gong, R. Wu and Y. Zhang, Journal of computational chemistry, 2015, 36, 2228–2235.

48 M. Q. Fatmi, T. S. Hofer, B. R. Randolf and B. M. Rode, The Journal of chemical physics, 2005, 123, 054514.

49 R. Mehmood and H. J. Kulik, Journal of Chemical Theory and Computation, 2020, 16, 3121–3134.

50 Q. Cui, T. Pal and L. Xie, The Journal of Physical Chemistry B, 2021, 125, 689–702.

51 H. M. Senn and W. Thiel, Angewandte Chemie International Edition, 2009, 48, 1198–1229.

52 P. Vidossich and A. Magistrato, Biomolecules, 2014, 4, 616–645.

53 J. Behler and M. Parrinello, Physical review letters, 2007, 98, 146401.

54 J. S. Smith, O. Isayev and A. E. Roitberg, Chemical science, 2017, 8, 3192–3203.

55 V. L. Deringer and G. Csányi, Physical Review B, 2017, 95, 094203.

56 L. Zhang, J. Han, H. Wang, W. Saidi, R. Car and E. Weinan, Advances in Neural Information Processing Systems, 2018, pp. 4436–4446.

57 K. T. Schütt, H. E. Sauceda, P.-J. Kindermans, A. Tkatchenko and K.-R. Müller, The Journal of Chemical Physics, 2018, 148, 241722.

58 J. Vandermause, S. B. Torrisi, S. Batzner, Y. Xie, L. Sun, A. M. Kolpak and B. Kozinsky, npj Computational Materials, 2020, 6, 1–11.

59 T. W. Ko, J. A. Finkler, S. Goedecker and J. Behler, Accounts of Chemical Research, 2021, 54, 808–817.

60 C. Devereux, J. S. Smith, K. K. Huddleston, K. Barros, R. Zubatyuk, O. Isayev and A. E. Roitberg, Journal of Chemical Theory and Computation, 2020, 16, 4192–4202.

61 A. Hofstetter, L. Böselt and S. Riniker, Physical Chemistry Chemical Physics, 2022, 24, 22497–22512.

62 L. Böselt, M. ThuÍĹrlemann and S. Riniker, Journal of Chemical Theory and Computation, 2021, 17, 2641–2658.

63 C. Qu, Q. Yu, R. Conte, P. L. Houston, A. Nandi and J. M. Bowman, arXiv preprint arXiv:2206.04254, 2022.

64 L. Zhang, H. Wang, R. Car and E. Weinan, Physical Review Letters, 2021, 126, 236001.

65 M. F. C. Andrade, H.-Y. Ko, L. Zhang, R. Car and A. Selloni, Chemical Science, 2020, 11, 2335–2341.

66 F.-Z. Dai, B. Wen, Y. Sun, H. Xiang and Y. Zhou, Journal of Materials Science & Technology, 2020, 43, 168–174.

67 J. Huang, S. Rauscher, G. Nawrocki, T. Ran, M. Feig, B. L. De Groot, H. Grubmüller and A. D. MacKerell, Nature methods, 2017, 14, 71–73.

68 F. Neese, Wiley Interdisciplinary Reviews: Computational Molecular Science, 2012, 2, 73–78.

69 B. R. Brooks, C. L. Brooks III, A. D. Mackerell Jr, L. Nilsson, R. J. Petrella, B. Roux, Y. Won, G. Archontis, C. Bartels, S. Boresch et al., Journal of computational chemistry, 2009, 30, 1545–1614.

70 J. Zeng, D. Zhang, D. Lu, P. Mo, Z. Li, Y. Chen, M. Rynik, L. Huang, Z. Li, S. Shi et al., The Journal of Chemical Physics, 2023, 159, 054801.

71 P. Eastman, J. Swails, J. D. Chodera, R. T. McGibbon, Y. Zhao, K. A. Beauchamp, L.-P. Wang, A. C. Simmonett, M. P. Harrigan, C. D. Stern et al., PLoS computational biology, 2017, 13, e1005659.

72 S. Jo, T. Kim, V. G. Iyer and W. Im, Journal of computational chemistry, 2008, 29, 1859–1865.

73 A. E. Eriksson, T. A. Jones and A. Liljas, Proteins: Structure, Function, and Bioinformatics, 1988, 4, 274–282.

74 Y. Xue, A. Liljas, B.-H. Jonsson and S. Lindskog, Proteins: Structure, Function, and Bioinformatics, 1993, 17, 93–106.

75 K. Kannan, M. Ramanadham and T. Jones, Annals of the New York Academy of Sciences, 1984, 429, 49–60.

76 D. Lu and G. A. Voth, Proteins: Structure, Function, and Bioinformatics, 1998, 33, 119–134.

77 D. C. Rees, M. Lewis and W. N. Lipscomb, Journal of molecular biology, 1983, 168, 367–387.

78 S. P. Dagnall, D. N. Hague and A. D. Towl, Journal of the Chemical Society, Faraday Transactions 2: Molecular and Chemical Physics, 1982, 78, 2161–2167.

79 E. Cauët, S. Bogatko, J. H. Weare, J. L. Fulton, G. K. Schenter and E. J. Bylaska, The Journal of chemical physics, 2010, 132, 194502.

80 J. Zeng, T. J. Giese, S. Ekesan and D. M. York, Journal of chemical theory and computation, 2021, 17, 6993–7009.

81 X. Pan, J. Yang, R. Van, E. Epifanovsky, J. Ho, J. Huang, J. Pu, Y. Mei, K. Nam and Y. Shao, Journal of Chemical Theory and Computation, 2021, 17, 5745–5758.

82 S.-L. J. Lahey and C. N. Rowley, Chemical science, 2020, 11, 2362–2368.

83 D. A. Rufa, H. E. B. Macdonald, J. Fass, M. Wieder, P. B. Grinaway, A. E. Roitberg, O. Isayev and J. D. Chodera, BioRxiv, 2020.

84 R. Galvelis, A. Varela-Rial, S. Doerr, R. Fino, P. Eastman, T. E. Markland, J. D. Chodera and G. De Fabritiis, Journal of Chemical Information and Modeling, 2023.

85 Y. Ding, K. Yu and J. Huang, Current Opinion in Structural Biology, 2023, 78, 102502.

86 Y. Zhang, H. Wang, W. Chen, J. Zeng, L. Zhang, H. Wang and E. Weinan, Computer Physics Communications, 2020, 253, 107206.

87 W. Maret, Biochemistry, 2004, 43, 3301–3309.

