## Supplemental Material for "DP/MM: A Hybrid Model for Zinc-Protein Interactions in Molecular Dynamics"

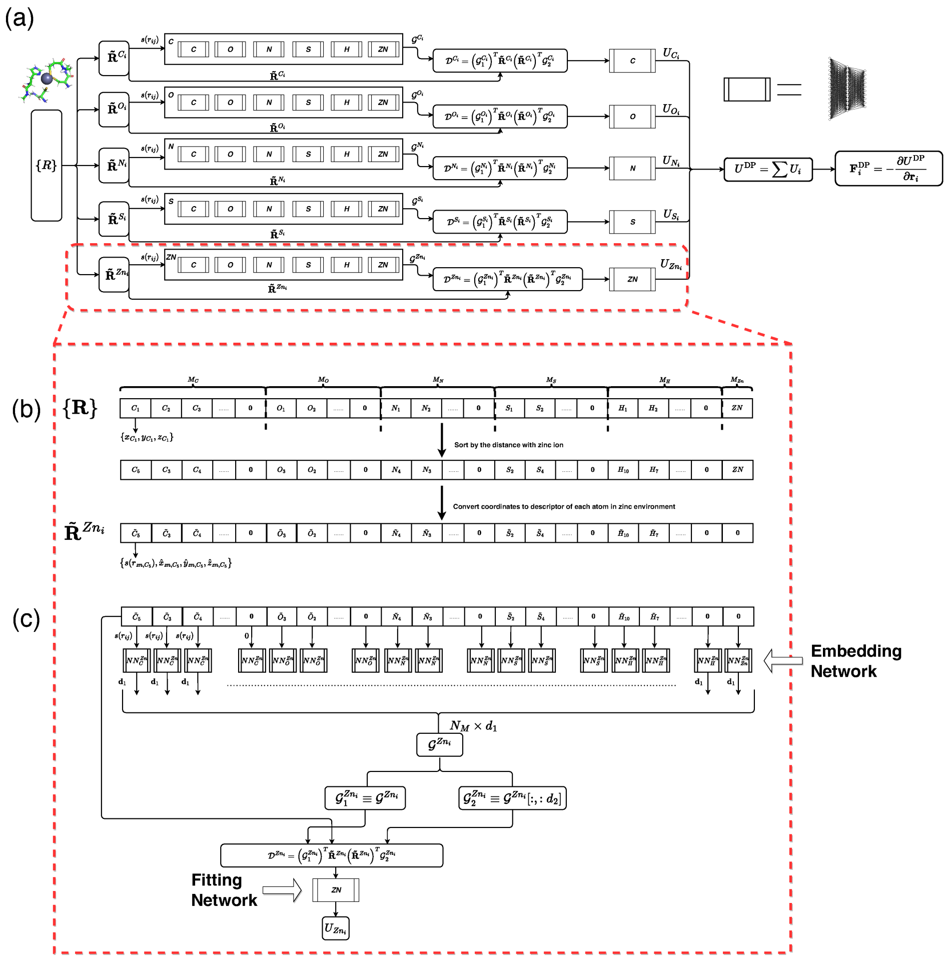


**Figure S1.** An overview of the neural network architecture for the DP model is depicted in (a). The coordination of zinc ion and its coordinated atoms in the lab frame is represented by $\{\mathbf{R}\}$. Firstly, the $\{\mathbf{R}\}$ is transformed to the relative coordinates $\{\tilde{\mathbf{R}}^{Zn_i}\}$ that utilized $\mathbf{r}^{Zn_i}$ as centered atom as (b) illustrated protocol. As the protocol shown in (b), the relative coordinates of $Zn_i$'s neighbors are sorted by distance to $Zn_i$ for permutation invariant. And the relative coordinates, e.g $x_{Zn_i,C_5}, y_{Zn_i,C_5}, z_{Zn_i,C_5}$, are converted to $s_{Zn_i,C_5}, \tilde{x}_{Zn_i,C_5}, \tilde{y}_{Zn_i,C_5}, \tilde{z}_{Zn_i,C_5}$ by the following equations:

$$s_{Zn_i,C_5} \equiv \frac{1}{|\mathbf{r}_{Zn_i,C_5}|}, \quad \tilde{x}_{Zn_i,C_5} \equiv x_{Zn_i,C_5} \cdot s_{Zn_i,C_5}, \quad \tilde{y}_{Zn_i,C_5} \equiv y_{Zn_i,C_5} \cdot s_{Zn_i,C_5}, \quad \tilde{z}_{Zn_i,C_5} \equiv z_{Zn_i,C_5} \cdot s_{Zn_i,C_5}. \tag{1}$$

Then, the "weights" factors $s(r_{ij})$ (inverse of the distance between the central atom and neighbor atom) are fed into the embedding network as the element type combination of the central atom and neighbor atom, (c). Note that the number of neighbor atoms of the central atom for each element is fixed in the DP model, i.e. carbon is 30, oxygen is 16, nitrogen is 24, hydrogen is 64, sulfur is 6 and zinc is 1. If the number of neighbor atoms is less than the fixed number, dummy atoms are padded up to the neighbor atoms list and the "weights" factors for this atom and other atoms are set to zero, which can be interpreted as the virtual atoms are infinite far away from the centered real atom. Finally, the output of embedding networks is concatenated together as $\mathcal{G}^{Zn_i}$, and multiplied with $\mathbf{R}^{\tilde{Z}n_i}$ and its transpose for rotation invariant. Feed the environment vectors $\mathcal{D}^{Zn_i}$ into the fully connected element-wised fitting networks, and the output of the fitting network is regarded as atomic energy contribution to the total energy of the system.

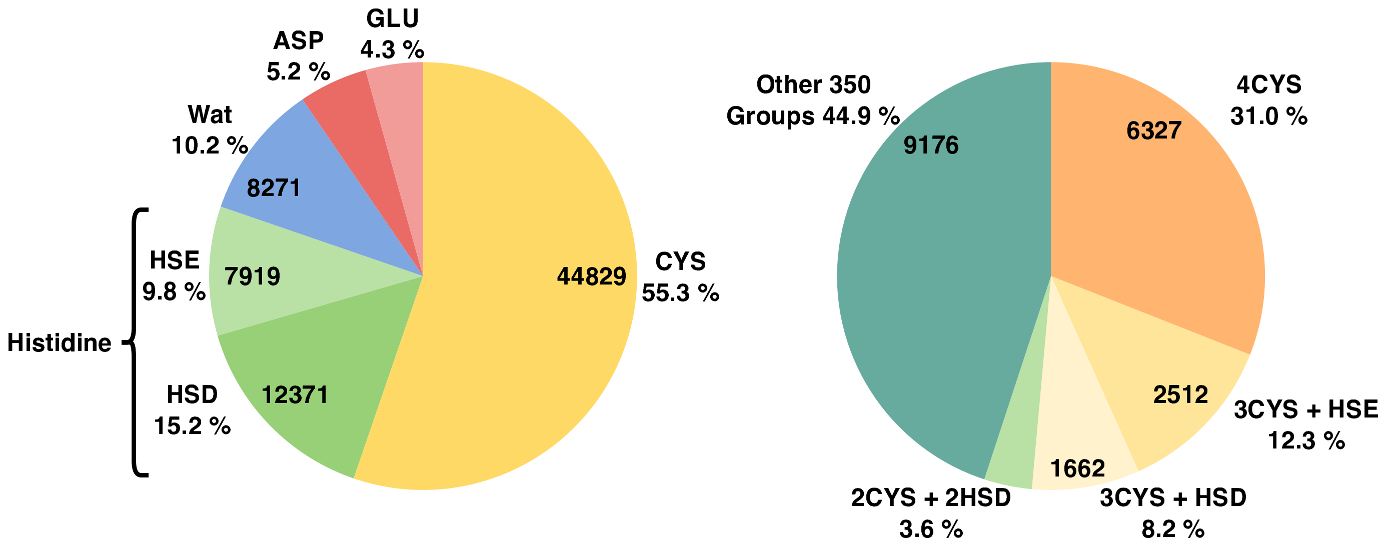


**Figure S2.** Pie chart of the occupancy of the selected residues (left) and coordination groups (right) in the crystal structure dataset only.

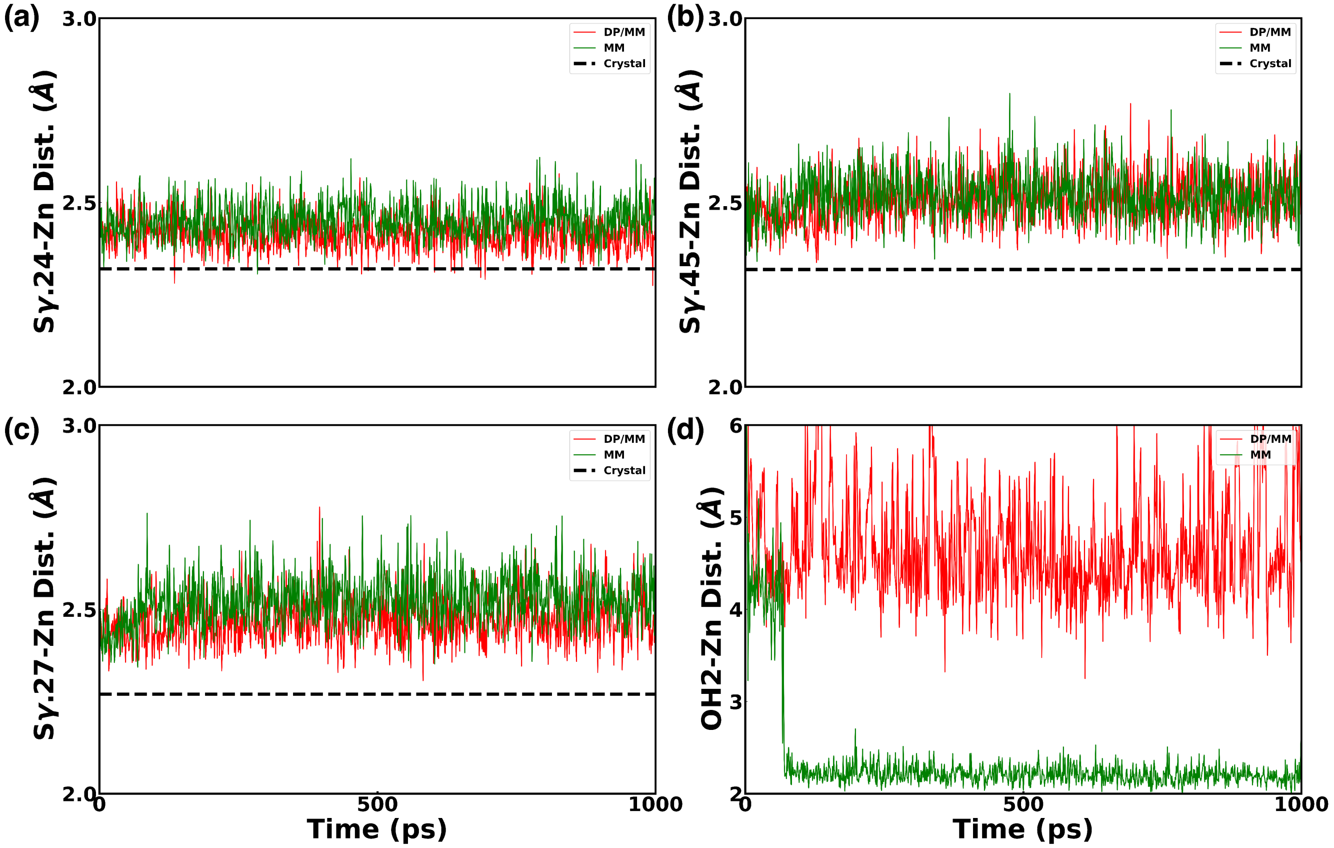


**Figure S3.** Distances evolution of three coordinated atoms S$\gamma$.24, 27, 45 with zinc ion in protein 6ZFV are illustrated in panels (a), (c) and (b) respectively. Besides, the distance of the nearest water molecule with zinc ion in DP/MM and MM simulations is also shown in panel (d).

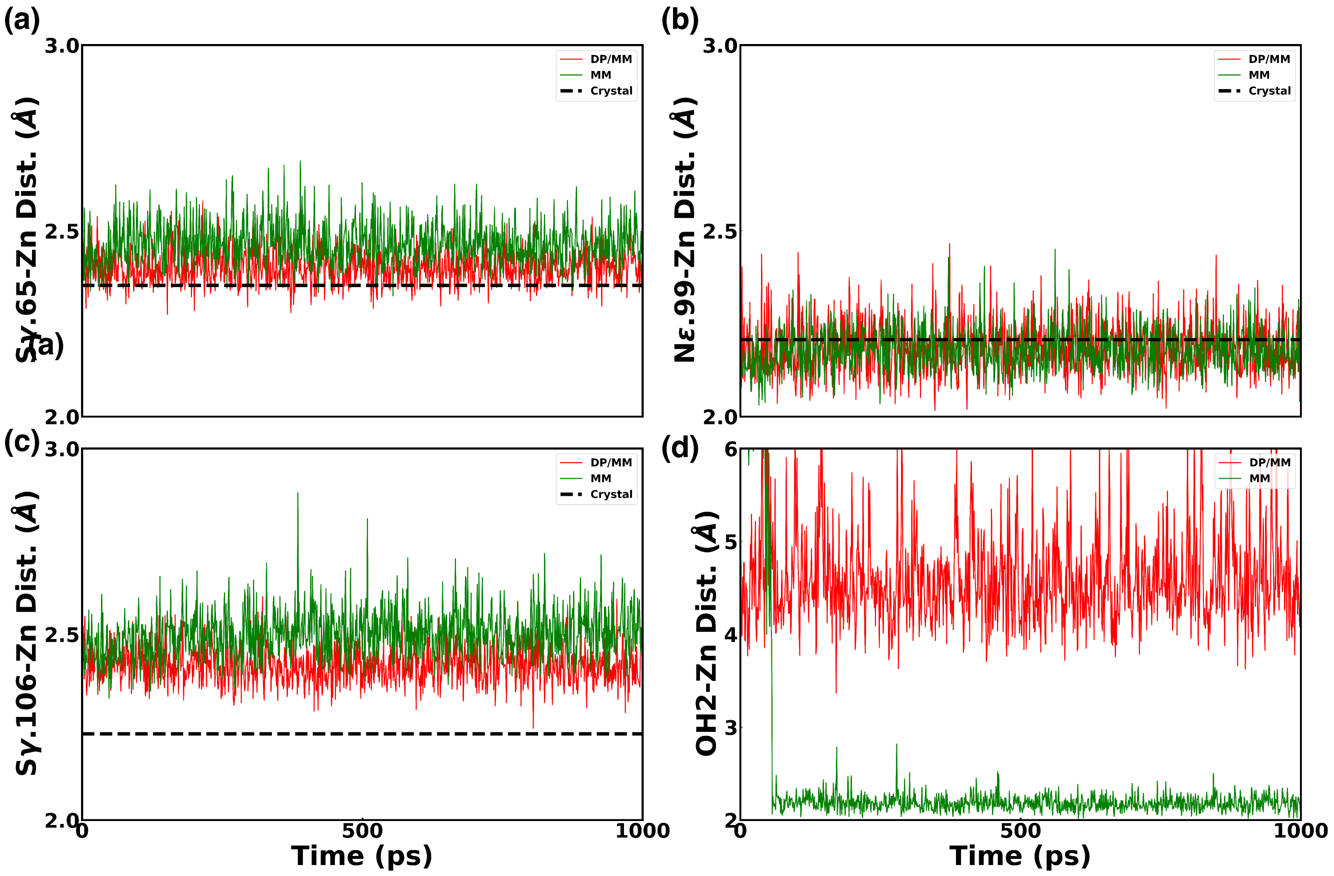


**Figure S4.** Distances evolution of the coordinated atoms S$\gamma$.65, 106 and N$\epsilon$.99 with zinc ion in protein 7EEZ are illustrated in panels (a) (c) and (b) respectively. The distance of the nearest water molecule with zinc ion in DP/MM and MM simulations is shown in panel (d).

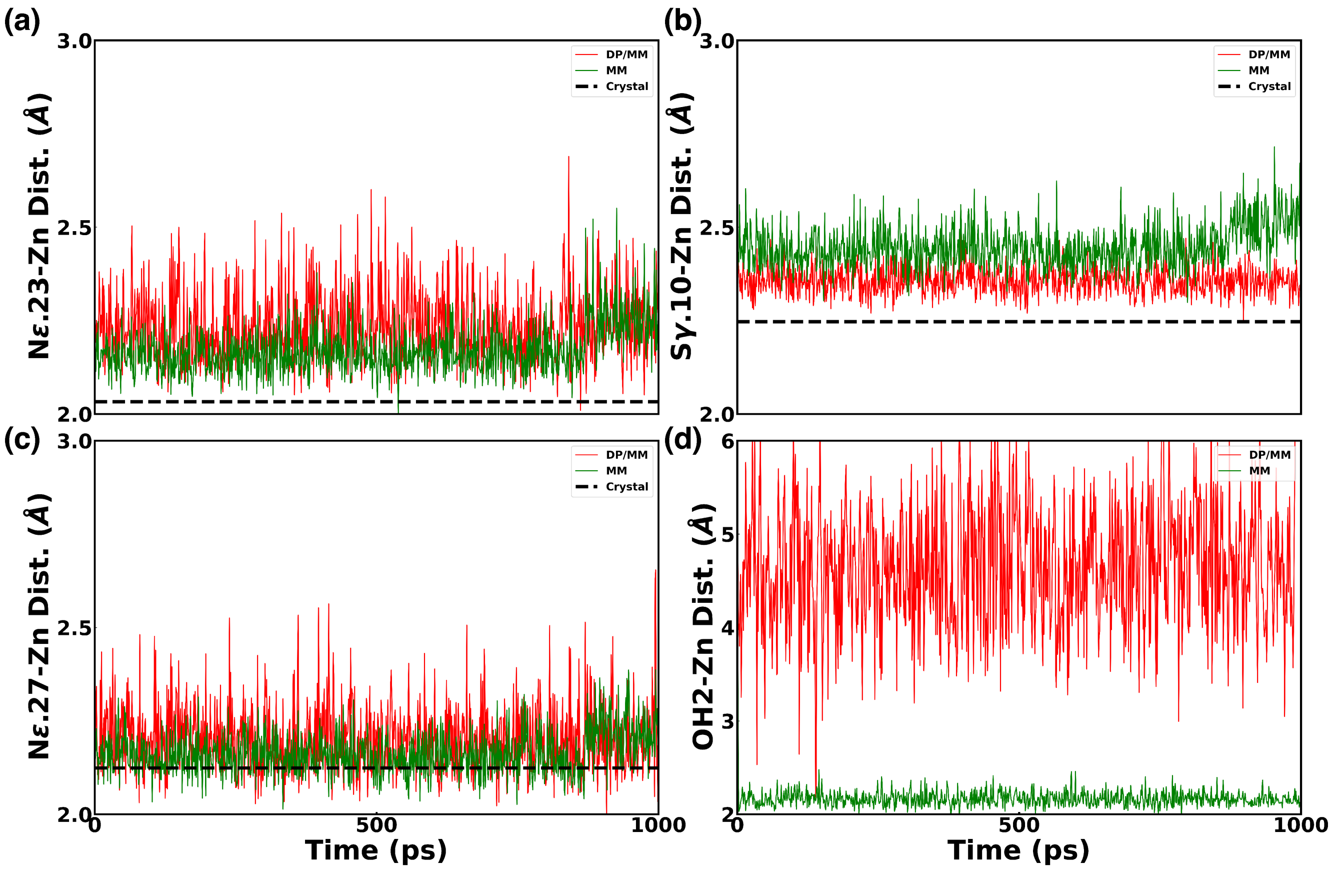


**Figure S5.** Distances evolution of the coordinated atoms N$\epsilon$.23, 27 and S$\gamma$.10 with zinc ion in protein 1SP2 are illustrated in (a), (c) and (b) respectively. The distance of the nearest water molecule with zinc ion in DP/MM and MM simulations is shown in panel (d).

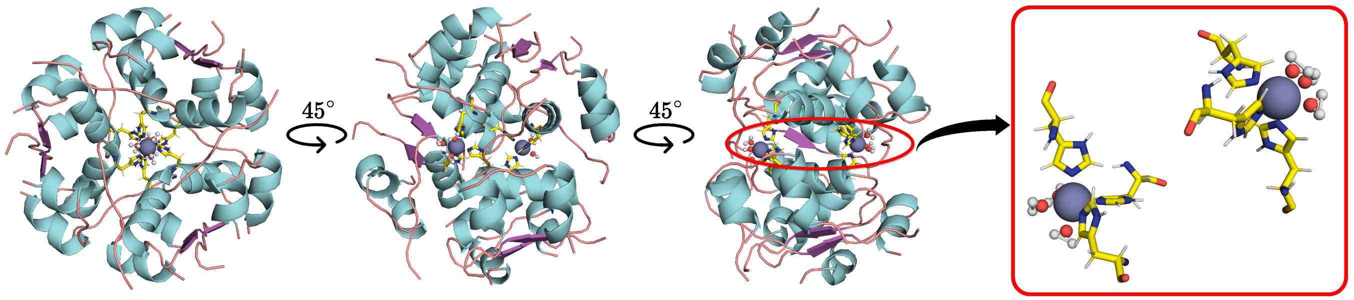


**Figure S6.** Crystal Structure details of 1MSO protein.

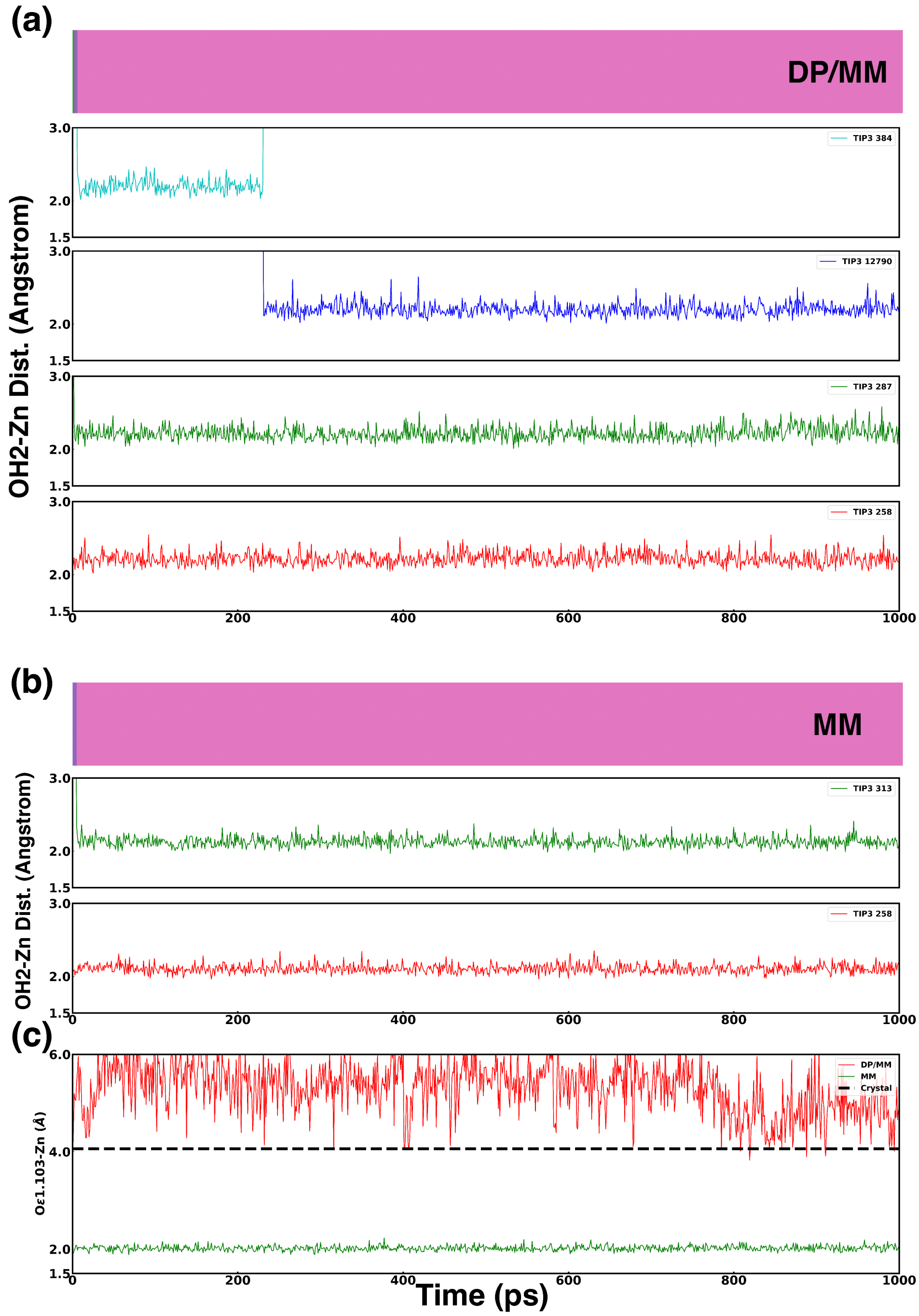


**Figure S7.** The time series of the distances between zinc and its coordinated water molecules in protein 1CA2 is illustrated in panels (a) (DP/MM) and (b) (MM). To show the major difference between DP/MM and MM simulation for 1CA2, the distances of zinc ion with the O$\epsilon$1 atom of Glu 103 are shown in panel (c)

**Table S1.** To validate the stability of the formed tetrahedral coordination geometry in longer DP/MM simulations. Two 10 ns DP/MM simulations were performed for 6ZFV and 7EEZ proteins. During these two 10 ns DP/MM simulations, the coordination number (CN) value for zinc ions was always 4. No additional water molecules were observed to be coordinated with zinc ions during the 10 ns DP/MM simulations. Besides, as the table shows, the distances of zinc ions and their coordinated atoms were stable in the 10 ns DP/MM simulations. Unit is angstrom.

| Protein | Dist. Type | MM (1 ns) | DP/MM (1 ns) | DP/MM (10 ns) | Crystal |
|---|---|---|---|---|---|
| 6ZFV | S$\gamma$.24-Zn | 2.452 (0.051) | 2.415 (0.047) | 2.438 (0.059) | 2.320 |
| | S$\gamma$.27-Zn | 2.525 (0.072) | 2.464 (0.061) | 2.452 (0.054) | 2.270 |
| | S$\gamma$.45-Zn | 2.521 (0.069) | 2.504 (0.066) | 2.472 (0.060) | 2.318 |
| | S$\gamma$.48-Zn | 2.536 (0.075) | 2.436 (0.046) | 2.428 (0.047) | 2.290 |
| 7EEZ | S$\gamma$.65-Zn | 2.466 (0.057) | 2.401 (0.044) | 2.401 (0.045) | 2.353 |
| | S$\gamma$.104-Zn | 2.539 (0.075) | 2.415 (0.048) | 2.394 (0.044) | 2.386 |
| | S$\gamma$.106-Zn | 2.502 (0.065) | 2.394 (0.043) | 2.415 (0.048) | 2.232 |
| | N$\epsilon$.99-Zn | 2.179 (0.058) | 2.181 (0.065) | 2.182 (0.068) | 2.207 |

**Table S2.** Distances of the coordinated atoms with two zinc ions in 1MSO protein are listed. Both two zinc ions and their coordinated residues were embedded in DP regions in DP/MM simulation. On average, N$\epsilon$-Zn distances in DP/MM simulation were similar to MM simulation results. OH2-Zn distances in DP/MM simulation were significantly larger ($\tilde{0}.07$ Å) than MM simulation results, and closer to the values in the crystal structure. The unit is in Angstrom.

| | MM | DP/MM | Crystal |
|---|---|---|---|
| N$\epsilon$.31-Zn.103 | **2.194** (0.058) | 2.201 (0.074) | 2.093 |
| N$\epsilon$.629-Zn.103 | **2.186** (0.057) | 2.206 (0.075) | 2.093 |
| N$\epsilon$.331-Zn.103 | **2.195** (0.060) | 2.201 (0.075) | 2.093 |
| OH2.736-Zn.103 | 2.113 (0.059) | **2.187** (0.085) | 2.204 |
| OH2.438-Zn.103 | 2.111 (0.058) | **2.185** (0.085) | 2.204 |
| OH2.140-Zn.103 | 2.110 (0.058) | **2.192** (0.087) | 2.204 |
| N$\epsilon$.680-Zn.104 | **2.189** (0.058) | 2.208 (0.078) | 2.102 |
| N$\epsilon$.82-Zn.104 | **2.192** (0.058) | 2.201 (0.075) | 2.102 |
| N$\epsilon$.382-Zn.104 | **2.192** (0.058) | 2.201 (0.075) | 2.102 |
| OH2.542-Zn.104 | 2.109 (0.058) | **2.192** (0.088) | 2.234 |
| OH2.840-Zn.104 | 2.110 (0.058) | **2.190** (0.089) | 2.234 |
| OH2.244-Zn.104 | 2.112 (0.058) | **2.191** (0.088) | 2.234 |

**Table S3.** Simulation recipes for each zinc-containing system.

| System | Num. of Atoms | Box (Å) | Temperature (K) |
|---|---|---|---|
| 6ZFV | 18254 | $64 \times 64 \times 64$ | 300 |
| 7EEZ | 41156 | $76 \times 76 \times 76$ | 300 |
| 1SP2 | 17317 | $57 \times 57 \times 57$ | 300 |
| 1CA2 | 48254 | $80 \times 80 \times 80$ | 300 |
| 1MSO | 39979 | $75 \times 75 \times 75$ | 300 |
| 5CPA | 61540 | $87 \times 87 \times 87$ | 300 |
| Zinc-Water | 1717 | $26 \times 26 \times 26$ | 300 & 350 |
